# Biosensor-Assisted Laboratory Evolution of Malonyl-CoA production in *Saccharomyces cerevisiae*

**DOI:** 10.1101/2023.07.16.549225

**Authors:** Raphael Ferreira, Karl Alex Hedin, Jens Nielsen, Florian David

**Author notes:** Corresponding author: Florian David, PhD Dr. Tech. Contributed equally (Raphael Ferreira, Karl Alex Hedin).

## Abstract

The production of bio-based chemicals and fuels through microbial engineering offers a promising and sustainable alternative to petroleum-based fuels and chemicals, with the potential for scalability. However, engineering microbes and continuously evolving them to enhance the production of industrially relevant products is a complex and challenging task, requiring precise selection of genetic traits to achieve desired outcomes. In this study, we report the development of a novel counter-selectable growth-sensitive malonyl-CoA platform strain by coupling the malonyl-CoA repressor FapR from *Bacillus subtilis* to essential gene promoters involved in glucose growth and the plasma membrane arginine permease. This platform strain was then coupled with a CRISPR-dCas9 guide-RNA (gRNA) library, which after multiple rounds of dilutions and library sequencing, resulted in the enrichment for gRNAs that increased fluxes towards malonyl-CoA. The enriched gRNAs were validated for their effects on growth enhancement, gene regulation, and the production of an industrially relevant malonyl-CoA product, namely 3-hydroxypropionic acid. This study highlights an innovative approach to microbial engineering and opens up avenues for further exploration in the field of laboratory continuous evolution.

## 1. Introduction

The advent of metabolic engineering of microbial cell factories for the production of industrially relevant compounds has been offered as one potential alternative to palliate the harmful effects of non-sustainable petrol based processes. Extensive genetic engineering of these microorganisms is required to reach economically viable production titers, rates and yields (TRY)^1^. An important strategy in microbial metabolic engineering is directing the central carbon metabolism towards the supply of metabolic precursors required for the synthesis of the compound of interest. For example, rewiring metabolism towards increased supply of malonyl-CoA is beneficial for the production of fatty-acids and derived products including biofuels, biopharmaceuticals such as phloroglucinol, or bioplastic such as 3-hydroxypropionic acid (3-HP)^2^. Malonyl-CoA is synthesized from acetyl-CoA by the enzyme acetyl-CoA carboxylase (Acc1) and is one of the endogenous precursors for iterative elongation of acyl chains^3,4^. This enzyme is a major regulation step in fatty acid biosynthesis, tightly regulated at the transcriptional and protein-level^4^. Expression of a double mutant *ACC1\*\** (*ACC1*^*S659A, S1157A*^), removing two phosphorylation sites, resulted in a constitutively active version of the enzyme^5–7^ and higher flux towards malonyl-CoA and derived products^8,9^. Several engineering strategies have been applied to enhance metabolic fluxes towards malonyl-CoA, mainly in *Escherichia coli* and *Saccharomyces cerevisiae*^8^. Other strategies have been relying on transcription factor-based biosensors, which offer novel high-throughput screening techniques to find high performing strains, ultimately allowing for a significant reduction of screening time. For example, in *E. coli*, Yang and colleagues recently applied a synthetic small regulatory RNA library coupled with a malonyl-CoA biosensor to identify gene knockdown targets enhancing the malonyl-CoA level^10^. Additionally, we recently applied a similar strategy where we combined a malonyl-CoA biosensor with a CRISPR guide-RNA (gRNA) library coupled to a dead-Cas9 fused to the tripartite VPR activator, allowing us to enrich for specific gRNAs which enhance fluxes towards malonyl-CoA^11–13^. However, our approach was constrained by the fact that it required the usage of fluorescent activated cell sorting (FACS) for high-throughput screening. In this context, one aspect of improving strain performance through biosensors is to associate them with an adaptive laboratory evolution (ALE) approach. For example, Leavitt and colleagues coupled a resistance driven biosensor and antimetabolites with ALE which ultimately resulted in the highest muconic acid titers in *S. cerevisiae* reported to date^14^.

Here we propose to apply a new type of optical detection for biosensor, namely optical cell density. The ease with measuring optical density (OD) allows for biosensor-assisted laboratory evolution (BALE) approach where high-performing stains, here strains displaying an enrichment of intracellular malonyl-CoA levels, will be selected through outgrowing other strains. Additionally, we implemented a canavanine counter-selection feature to our platform strain in order to reduce the risk of potential false-positives^15–17^.

## 2. Materials and Methods

### 2.1 Growth medium

Synthetic dextrose containing 6.7 g·L^-1^ of yeast nitrogen base without amino acids (DifcoLaboratories, Sparks, MD, USA), 0.77 g·L^-1^ of complete supplement mixture (CSM, without uracil and histidine) (MP Biomedicals, Solon, OH, USA), and 2% glucose were used on strains carrying *HIS3* and *URA3* marker(s). Strains carrying a KanMX resistance marker were selected on YPD plates containing 10 g·L^-1^ yeast extract, 20 g·L^-1^ casein peptone, 20 g·L^-1^ glucose, and 20 g·L^-1^ agar were supplemented with 200 mg·L^-1^ G418 (Formedium). All cultures for growth measurement were cultured in minimal medium containing 20 g·L^-1^ glucose, 5 g·L^-1^, (NH_4_)_2_SO_4_, 14.4 g·L^-1^ KH_2_PO_4_, 0.5 g·L^-1^ MgSO4·7H_2_O, complemented with histidine and/or uracil. After sterilization, 2 mL·L^-1^ trace element solution and 1 mL·L^-1^ of vitamin solution were added.

### 2.2 Strain and plasmid construction

Oligonucleotide and gRNA sequences were ordered from Eurofins and integrated DNA Technologies, IDT and are listed in Supplementary information Tables S1. Phusion high-fidelity DNA polymerase (Thermo Scientific, Waltham, MA, USA) and PrimeSTAR HS (TaKaRa) were used for all PCR reactions. All plasmids and integration cassettes were cloned using Gibson Assembly. FapR and GFP were amplified from pFDA09 and pRS413TEF-GFP respectively and assembled with the integration plasmid p395 and digested with NotI prior integration. The modified TEF1-BS123 promoter was amplified from pFDA10 with respectively overlap to the genome.

*Saccharomyces cerevisiae* CEN.PK113-11C (*MAT*a *his3*Δ*1 ura 3-52 MAL2-8c SUC2*) was obtained from P. Kötter, University of Frankfurt, Germany. The strains used in this study are listed in Supplementary information Table S1. *S. cerevisiae* transformations were performed with the high-efficiency yeast transformation using the LiAc/SS carrier DNA/PEG method^18^. CRISPR techniques were used for integration of all modified *TEF1-*BS123 promoters and yielded strains AHP01-AHP04 and MCP01. Benchling CRISPR tool generated all the gRNA sequences. Integration of *TEF1*-BS123 was confirmed with colony-PCR using DreamTaq (Thermo Scientific, Waltham, MA, USA). Genomic DNA extraction was performed by boiling biomass in 15 μL of 20 mM NaOH for 15 min and centrifuged at max speed for 15 seconds.

### 2.3 Real-time growth monitoring and fluorescence measurements

Real-time OD_600_ measurement was obtained every 30 min for approximately 72 hours with *Growth profiler 960*. The cultures were incubated into 250 μL minimal medium, in a PS 96-half deep well microtiter plate with air-penetrable lid (Duetz System, Kuhner Shaker). The cultivation was done at 250 rpm in 30°C and with an initial OD_600_ of 0.05. Real-time monitoring of GFP expression was obtained every 15 min for approximately 24 hours with a BioLector ® (m2p-labs GmbH, Baesweiler, Germany). The cultures were incubated into 1 mL minimal medium, in a FlowerPlates ®. The cultivation was done at 250 rpm in 30°C and with an initial OD_600_ of 0.05. GFP expression levels were measured by the ratio of green fluorescence/biomass.

### 2.4 Counter-selection

L-canavanine was purchased from Sigma Aldrich. Three different canavanine concentrations (0, 3 and 10 μg·mL^-1^) were evaluated for the platform strain MCP01 with and without FapR. The most suitable concentration was evaluated based on the highest growth ratio between the strains with and without FapR. To evaluate the growth performance after the canavanine treatment the platform strain was first treated with 3 ng·μL^-1^ canavanine for 24 hours. 24 hours were chosen due to match the condition planned to use for the ALE. The cells were watched twice with minimal medium. Cells were later cultivated for a total of 108 hours.

### 2.5 CRISPR-dCas9 gRNA library

The CRISPR-dCas9 gRNA library was ordered from Twist Bioscience (San Francisco, CA, USA) and amplified with gRNAextender-F and gRNAextender-R. The amplified library was cloned into pDTU-113 and transformed directly into the MCP01 platform strains following Benatuil and colleagues high-efficiency electroporation protocol^19^. For optimized cloning, 1:3 vector-insertion ratio was used; 300 ng of digested vector and 900 ng of PCR amplified gRNA library. To increase the coverage of gRNAs the transformation was duplicated. The library was grown on appropriate selective plates and around 10000 yeast colonies (corresponding to three times the library size) were pooled together. The colonies were resuspended in minimal medium and stored in -80°C.

### 2.6 ALE (Adaptive Laboratory Evolution)

Triplicates of MCP01 couple with the CRISPR-dCas9 gRNA library were cultivated in 250 μL minimal medium, with anhydrotetracycline (aTc) (1000x) added, in a PS 96-half deep well microtiter plate with air-penetrable lid. The cultures were continuously transferred into fresh medium every 12 hours for a period of 108 hours and continued for a total period of 176 hours to measure the stability of the strain. After the second transfer (t=24h), we expose the cells with canavanine for 24 hours. The cells exposed to canavanine were washed twice and grown for 36 hours to restore growth (Figure S2). An initial OD_600_ of 0.05 was used after each transfer, except after addition of canavanine, where we used an OD_600_ of 0.1.

### 2.7 Next-generation sequencing analysis

After each transfer, 100 μL of culture were centrifuged at 12,000 g and the genomic DNA was extracted using Lõoke and colleagues genomic extraction protocol^20^. The gRNA libraries were amplified and prepared based on Illumina DNA Nextera Sequencing as described in Lee and colleagues^21^. All next-generation sequencing was performed on a MiSeq Benchtop Sequencer (Illumina, San Diego, CA) and a custom R script was used to analyze the Fastq sequences. The sequence was probed between the end of RPR1p (5’-CGATTGGCAG) and beginning of RPR1t (5’-GTTTTAGAGC) and matched to the gRNA library. The abundance, i.e. the number of times a gRNA is matched to the gRNA library, was quantified and normalized by the total number of NGS read counts for each run. The most enriched gRNAs were selected based on three criteria *(i)* taking the top 100 highest base 2 log ratio of the normalized gRNAs hits over the NGS from the preculture for each replicate at the time point 108 hours. The significantly enriched gRNAs, i.e. common occurring among all the replicate’s top 100, were selected; *(ii)* receiving the top 20 highest gradual distribution enrichment of the normalized distributed gRNAs in MCP01F; *(iii)* taking the top 20 gRNAs with highest difference between the distribution enrichment over time of MCP01F’s and MCP01C’s. The figures were plotted using the R library ggplot2 3.3.0.

### 2.8 3-Hydroxypropionic acid measurements

Triplicates of the CEN.PK113-11C carrying MCR expressing gRNAs were cultivated aerobically in 10 mL minimal medium, with aTc (1000x) added, in shake flasks with a shaking speed of 200 rpm. All culture samples were collected after 12 hours, centrifuged at 12,000g and the supernatants were diluted 1:5 with 0.5 mM H_2_SO_4_. A high-performance liquid chromatography (HPLC, Dionex UltiMate 3000; Thermo Fisher Scientific, Waltham, MA, USA), equipped with an Aminex HPX-87H (Bio-Rad, Hercules, USA) column at 65°C, with a mobile phase of 0.5 mM H2SO4 at a flow rate of 0.5 mL/ min for 35 min were used for measure the 3-HP concentration.

## 3. Results

### 3.1 Construction of a FapR-based growth-sensitive counter-selectable malonyl-CoA platform strain

Two parameters were considered to design a growth-sensitive malonyl-CoA platform strain:

*(i)* an intracellular sensor system able to detect perturbation in malonyl-CoA levels was incorporated; and *(ii)* desired sensitivity-response to malonyl-CoA was robustly coupled to growth, i.e. any perturbation leading to high-levels of malonyl-CoA would confer a selective growth advantage over cells with low-levels. We based our system on the previously characterized malonyl-CoA-sensitive transcription factor FapR derived from *Bacillus subtilis* and its binding sites integrated into a *TEF1* promoter (P_*TEF1*_-BS123)^16^. In this context, the repression is alleviated by increased levels of malonyl-CoA, which binds to FapR and prevents its binding to P_*TEF1*_-BS123. Here, we genomically integrated P_*TEF1*_-BS123 to replace the endogenous promoters of two essential genes for growth on glucose, namely acetyl-CoA synthase 2 (*ACS2*)^22^, and phosphoglycerate kinase 1 (*PGK1*)^23^ (Figure 1). These native promoters were selected due to their transcriptional activity being in the range of P_*TEF1*_-BS123, which ultimately leads to substantial repression upon the expression of FapR (Table S1)^16,24^. *ACS2* and *PGK1* encode for enzymes that catalyze reactions upstream of malonyl-CoA: conversion of acetate to acetyl-CoA for *ACS2*, and 3-phospho-D-glycerate to 3-phospho-D-glyceroyl phosphate for *PGK1*. In theory, any transcriptional perturbation leading to an increased malonyl-CoA level will ultimately alleviate the repression of these two genes and ultimately boost growth. Notably, expression of FapR in wild type (AHP01F) did not result in a significant change in growth rate (Figure S1). Genomic replacement of the *ACS2* and *PGK1* promoter (AHP04C strain) with P_TEF1_-BS123 did not lead to a significant change in growth (Figure S1, Table S2). Expression of FapR (AHP04F) significantly affected the growth rate (Figure S1, Table S2).

**Figure 1.**
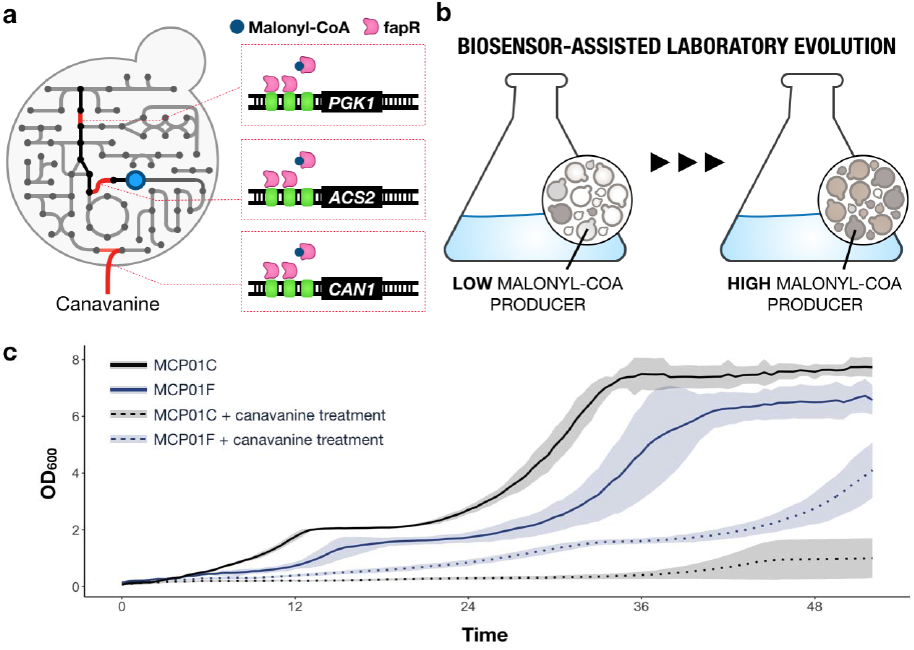
Construction of a FapR-based counter-selectable growth-sensitive malonyl-CoA platform strain. **a**. Graphical abstract of the engineered strain with PTEF1-BS123 integrated to *PGK1, ACS2*, and *CAN1* promoter loci. **b**. Graphical illustration of the scope of the BALE, achieving an enrichment of high malonyl-CoA producers. **c**. Growth performance of the engineered platform strain, MCP01, treated with 0 ng·μL^-1^ (plain) and 3 ng·μL^-1^ (dotted) of canavanine. The blue and gray lines describe the platform strain with and without FapR, respectively. Strains were grown in defined minimal medium with 20 g·L^-1^ glucose in triplicates.

Next, to remove potential false-positives, e.g. cells with a non-functional or lowly expressed FapR, we incorporated a counter-selection system to the platform strain. We explored the counter-selectable marker *CAN1*, which encodes for a plasma membrane arginine permease, and strains expressing this gene can be selected with canavanine, a toxic arginine analog^25^. FapR-repression of *CAN1* increases cell robustness by lowering canavanine toxic import, which ultimately allows removing cells carrying potential mutations in FapR that would die from high canavanine import. P_TEF1_-BS123 was subsequently integrated into the *CAN1* promoter region, yielding to a FapR-based counter-selectable growth-sensitive malonyl-CoA platform strain (MCP01). Optimal canavanine concentration, i.e. most significant growth difference between the strains with and without FapR, was found at 3 ng·μL^-1^ canavanine (Figure S2, Table S2) with a 4.4-fold higher OD_600_ in strains with FapR (Figure 1).

**Table 1.**
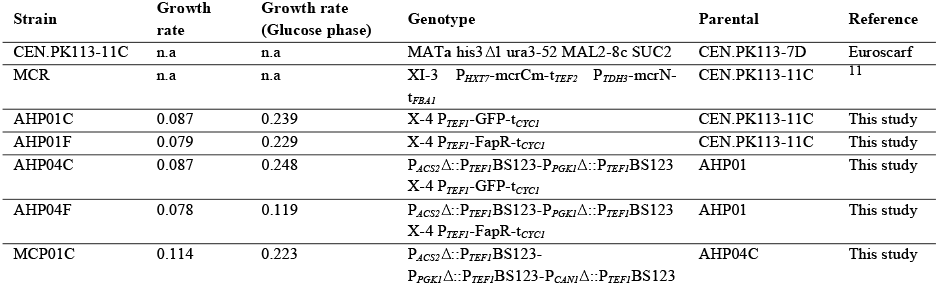
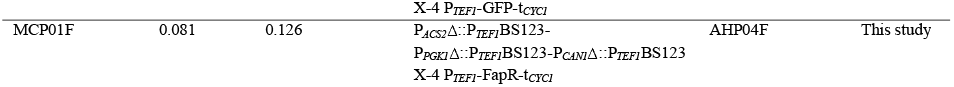
Strains used for this study.

### 3.2 Biosensor-assisted laboratory evolution of MCP01 combined with CRISPR/dCas9-based libraries

Next, we sought to exploit MCP01’s capability of sensing metabolic perturbation to identify candidate genes that enhance fluxes towards malonyl-CoA. In this context, CRISPR/Cas9-based technology offers a cost-efficient tool for targeting multiple specific genomic loci of interest and can conceivably be used to tune gene expression. We aimed to globally enhance metabolic fluxes towards malonyl-CoA by combining a dCas9-based gRNA library targeting 168 genes to our MCP01 platform strain. The 168 target genes were retrieved by performing a flux balance analysis (FBA) for two carbon sources, glucose and ethanol, where the specific growth rate (μmax) and acetyl-CoA and malonyl-CoA production were maximized as previously described^11^. The subsequent library was cloned under an anhydrotetracycline (aTc) inducible RNA polymerase III promoter into a centromeric plasmid encoding dCas9 coupled to a VPR activator domain through homologous recombination *in vivo* after transformation in MCP01C and MCP01F^13,26^. The yeast gRNA libraries were cultivated and diluted every 12 hours, corresponding to the window where the largest growth difference is observed between MCP01C and MCP01F. Canavanine was added to the ALE after the second dilution (t_24h_). Next-generation sequencing (NGS) was performed at every dilution stage and the gRNA distribution was determined (Figure 2). A distinct growth enrichment was seen in the engineered platform strain with the library compared to without, highlighting the potential of certain gRNAs to change gene regulation pattern leading to faster growth in MCP01 (Figure 2, S3). Here, we sought to generate three conditions to select gRNAs for further analysis: *(i)* MCP01F and MCP01C gRNAs distributions were normalized, and the base 2 log ratio over their respective preculture was determined. Here, we sorted the top 100 most enriched gRNAs after 108 hours in all three replicates of MCP01F. gRNAs significantly enriched in MCPO1F were selected and compared to MCP01C (Figure 2, Table S3-S4); *(ii)* next the normalized distributions of gRNAs in MCP01F were evaluated and the top 20 highest distribution enrichment over time were selected (Figure S4); *(iii)* additionally, the top 20 gRNAs with highest difference between the counts over time of MCP01F’s and MCP01C’s gRNAs were selected (Figure S4). Altogether, 8 different gRNAs fulfilled all three criteria (Figure 2, Table S5). None of the gRNAs displayed a distinct enrichment in the platform strain without canavanine treatment (Figue S5).

**Figure 2.**
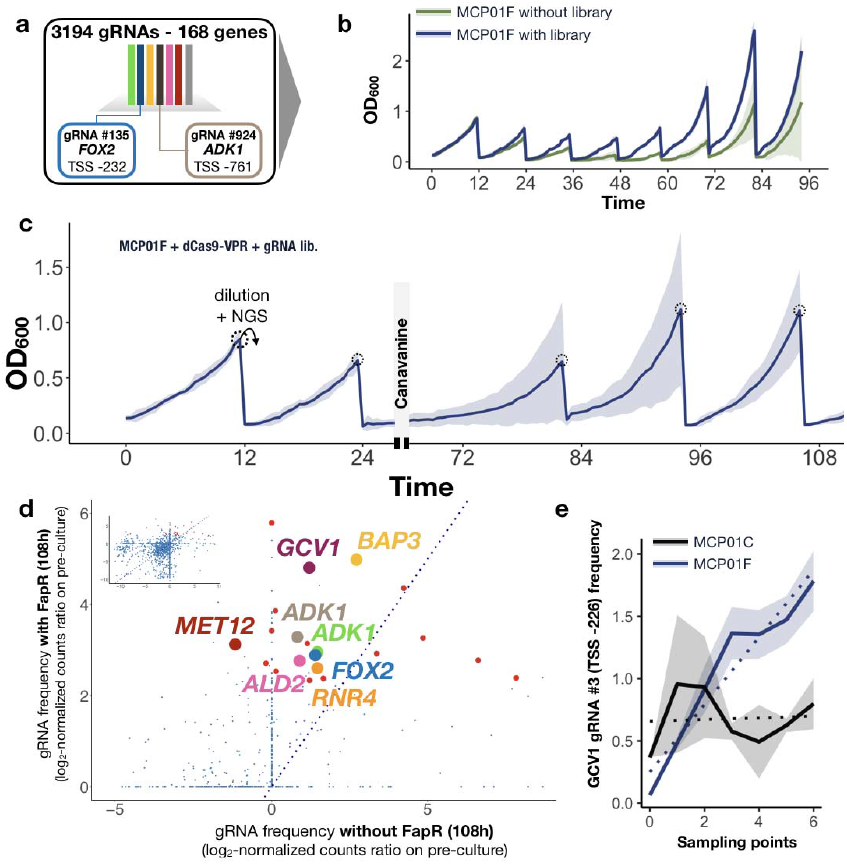
Biosensor-assisted laboratory evolution combined with CRISPR/dCas9-based libraries. **a**. A gRNA library of 3194 gRNAs designed to target 168 genes is coupled to an activator CRISPR/dCas9-VPR. Individual yeast (MCP01 background) expressing a gRNA whose transcriptional perturbation leads to enhanced levels of malonyl-CoA possess a growth advantage over the rest of the yeast culture and ultimately increase their frequency over time. **b**. Growth comparison of the engineered platform strain MCP01F with and without library, diluted every 12 hours **c**. Growth profile of the ALE. Cells were grown in 2% glucose minimal media and diluted every 12 hours. After 24 hours, i.e. second dilution point, cells were exposed to canavanine treatment. **d**. Log_2_ fold-change gRNAs at 12h over the initial library (pre-cultured) of MCP01F on the y axis and MCP01C on the x axis. Shown in red the enriched gRNAs that were tested for 3-HP production **e**. Example for frequency of the GCV1 gRNA #3 (TSS-226) evolving over time for each sample point comparing gRNA expressed in MCP01F and MCP01C. Content shown as mean of frequency ± SD of three biological replicates.

The stability of FapR and the strength of the counter-selection system to remove potential false-positives were evaluated by sequencing the FapR region after the last transfer for both cultures treated with and without canavanine. Cultures treated with canavanine at 24 and 108 hours, were grown for 176 hours in total, which corresponds to 8 transfers (Figure S3). Compared to 96 hours for the cultivation without canavanine added (Figure S3). Here we notice a lower degree of deviation in the culture not treated, displaying 46 % mutation compared to 12.5 % (Figure S6).

### 3.3 Enriched gRNAs fine-tune the expression of their targeted genes and enhance both growth fitness and production of a malonyl-CoA-derived product

We sought to individually assess the performance of the significantly enriched gRNAs for *(i)* growth fitness, i.e. growth rate improvement over the control (no gRNA expressed) in MCP01F and MCP01C; *(ii)* transcriptional regulation, i.e. fine-tuning properties of expressing the gRNAs on their respective targeted promoters; and *(iii)* enhancing fluxes towards malonyl-CoA pools, i.e. quantification of malonyl-CoA-derived product titers, here 3-hydroxypropionic acid (3-HP).

Growth fitness was evaluated by expressing each gRNA in MCP01F compared to the control, i.e. no gRNA expressed, as well as in MCP01C to assess whether the gRNAs had a general effect on growth (Figure 3, Table S7). Expression of ADK1 gRNA #15 (TSS -761), BAP3 gRNA #7 (TSS -47), FOX2 gRNA #17 (TSS -390), GCV1 gRNA #3 (TSS -226), MPC3 #5 (TSS -146), and RNR4 gRNA #12 (TSS -497) led to a growth increase compared to the strain with no gRNA expressed. Notably, through the expression of the gRNAs, the long lag phase displayed in MCP01 (Figure 1, 3) seemed to be fully recovered. Expression of the significantly enriched gRNAs in MCP01C resulted in a general growth increase for BAP3 gRNA #7 (TSS -47) and FOX2 gRNA #17 (TSS -390) (Figure S7). All gRNAs displayed a significant growth rate increase, except MPC3 #5 (TSS -146), p-value < 0.05. Likewise, an increased growth rate was displayed for BAP3 gRNA #7 (TSS -47), FOX2 gRNA #5 (TSS -323), MPC3 #12 (TSS -146), and RNR4 gRNA #12 (TSS -497) screened in MCP01C (Figure S7).

**Figure 3.**
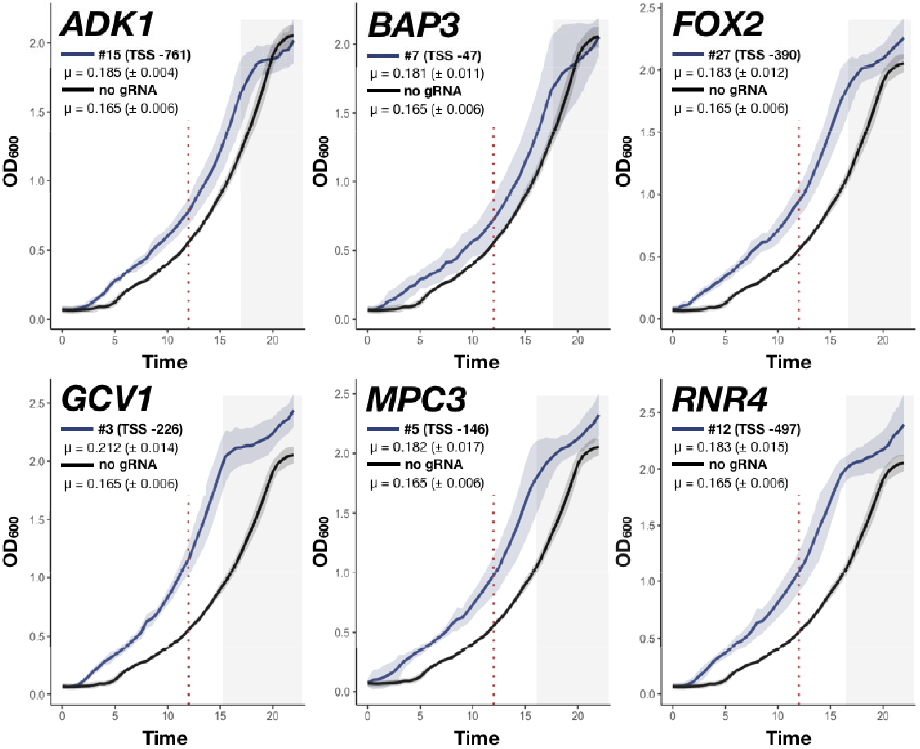
Characterization of the enriched gRNAs on growth fitness. The content shown as mean ± SD of three biological replicates and two technical replicates were monitored with a Growth Profiler 960. The growth rate was calculated between the start of the exponential phase to the 12-hour mark for all replicates ± SD. Outliers were removed (initial OD_600_ > 0.15).

The transcriptional regulation of the selected gRNAs was evaluated by coupling their targeted promoter to GFP. Here, the fluorescence levels are used as a proxy to assess the levels of transcriptional interference carried out by the gRNA:dCas9-VPR (Figure 4, Table S8). MPC3 gRNA #5 (TSS -146) and RNR4 gRNA #12 (TSS -497) show an upregulation pattern, while ADK1 gRNA #15 (TSS -761) and FOX2 gRNA #17 (TSS -390) show a down-regulating pattern. Notably, BAP3 gRNA #7 (TSS -47) and GCV1 gRNA #3 (TSS -226) displayed no significant change in fluorescence.

**Figure 4.**
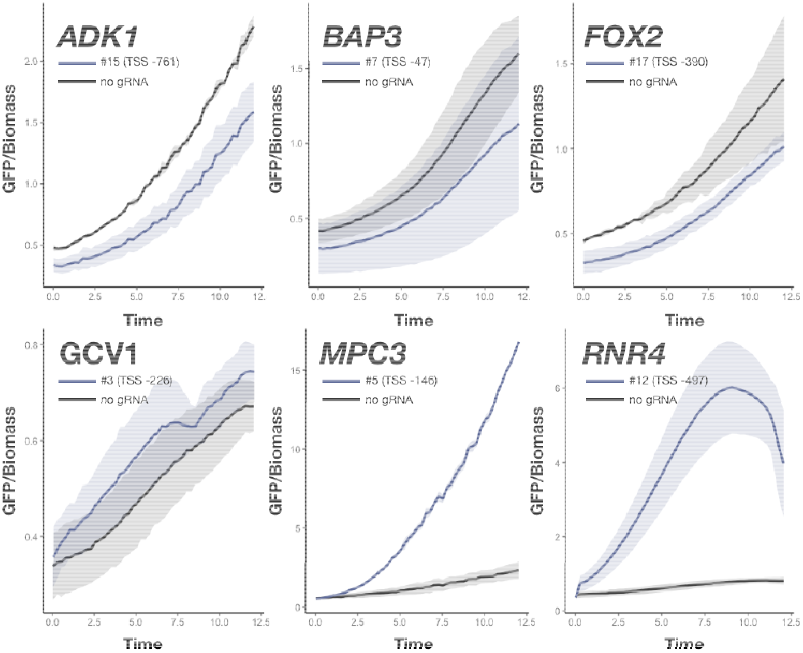
Characterization of the enriched gRNAs on transcriptional regulation. Content shown as mean ± SD of triplicates. Mean fluorescence intensities (GFP/OD) were obtained from three biological replicates ± S.D. monitored with a BioLector. Content shown as mean ± SD of three biological replicates and two technical replicates.

Finally, we evaluated the enhancing fluxes towards malonyl-CoA by producing 3-HP. The levels of 3-HP were validated through expressing the bifunctional malonyl-CoA reductase, *mcr* gene from *Chloroflexus aurantiacus*^27^. Here, we compared the performance upon expression of the selected gRNAs compared to the control (no gRNA expressed) after 12 hours cultivation, reflecting the dilution time point for the ALE (Figure 5, Table S5). Expression of ADK1 gRNA #15 (−761), BAP3 gRNA #7 (TSS -47), GCV1 gRNA #3 (TSS -226) and MPC3 #5 (TSS -146) resulted in significantly higher 3-HP yields compared to the control (no gRNA expressed).

**Figure 5.**
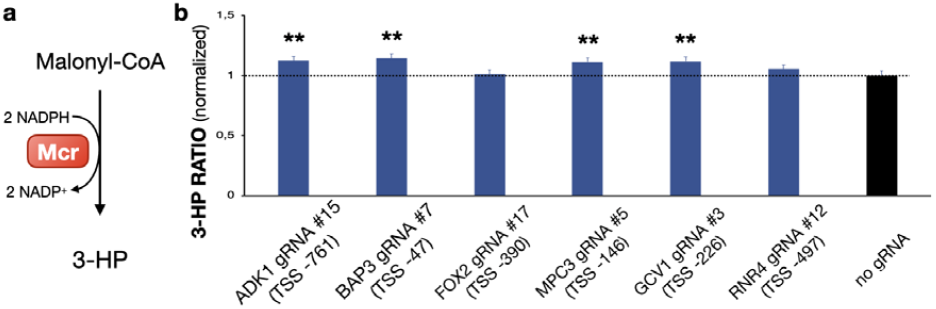
Characterization of the enriched gRNAs on the production of a malonyl-CoA derived product. **a**. Schematic of Malonyl-CoA conversion to 3-HP catalyzed by the bifunctional malonyl-CoA reductase Mcr from *Chloroflexus aurantiacus*. **b**. 3-HP production coupled to selected gRNAs. All gRNAs were normalized to OD_600_ and the control (no gRNA expressed). Content shown as mean ± SD of triplicates. \*\**p* value < 0.01 (Student’s *t* test: one-tailed, two-sample equal variance).

## 4. Discussion

Our BALE approach offers a cost-effective way to screen for beneficial setups, without requiring the need of FACS. The incorporation counterselecting system through a FapR-repression of *CAN1* ultimately allowed the removal of cells carrying potential mutations. Indeed, we observed several mutations occuring in FapR coding sequence after eight dilutions. Since we also observed mutations as well in the counterselecting platform, though to a lesser extent, more optimization might be required, e.g. using different essential genes. Additionally, while we validated our platform strain using CRISPR dCas9-based strategies we believe that other approaches such as mutagenized libraries or chemical drugs could be applied with the MCP01 platform strain.

We aimed to exploit the versatility of dCas9-VPR to rewire metabolic fluxes towards malonyl-CoA, by fine-tuning their expression level. Applying such a strategy has allowed us to retrieve new gene targets for metabolic engineering of *S. cerevisiae* for the production of malonyl-CoA derived products. CRISPR has now become an established tool for engineering microbial cell factories, and the field is observing more complex organisms, such as non-conventional yeasts, being successfully engineered with this technology^28–31^. As with our previous study^11^, ADK1 gRNA #15 (−761) showed the most significant results in the three assessed criterias. While regulating *ADK1* might have impacted ATP levels, which is used in the biosynthesis of malonyl-CoA from acetyl-CoA, upon closer inspection the gRNA also binds in the middle of *HTA1* coding sequence. *HTA1* encodes one of the major histone proteins and the gRNA binding might have reduced *HTA1* mRNA levels by blocking RNA polymerase II and thus potentially changed chromatin structures and consequently the expression levels of other genes. MPC3 #5 (TSS -146) shows an upregulating effect on *MPC3*, which encodes for a subunit of the mitochondrial pyruvate carrier (MPC). Theoretically, decreasing import of pyruvate into the mitochondria would have allowed more pyruvate to be converted to acetyl-CoA through the pyruvate decarboxylase (PDC) complex. However, Kildegaard and colleagues revealed through a transcriptome analysis of several 3-HP-producing *S. cerevisiae* strains that *MPC2*, which encodes for the subunit expressed during glucose fermentation, is upregulated^9^. Additionally, Mpc1 which dimerizes with Mpc3 during respiration has been characterized as a more efficient importer compared to its Mpc1/Mpc2 homolog^32^. This ultimately indicates a potential need for mitochondrial import of pyruvate for the production of 3-HP. Interestingly, while both NADPH/NADP+ and NADH/NAD+ ratios increase with the production of 3-HP, the absolute levels of NADH and NAD+ are drastically increased compared to NADPH and NADP+. Targeting *GCV1* showed increased levels of 3-HP and growth. *GCV1* encodes for a T subunit of the mitochondrial glycine decarboxylase complex which cleaves glycine into 5,10-Methylenetetrahydrofolate, a precursor for purines^33^. This compound can also be converted to tetrahydrofolate by the multifunctional mitochondrial C1-tetrahydrofolate synthase (Mis1), whose reactions involve NADPH and ATP cofactors. RNR4 gRNA #12 (TSS -497) also significantly increased growth rate, even though not changing 3-HP yields. *RNR4* encodes for a ribonucleotide-diphosphate reductase subunit in the purine biosynthesis pathway^34,35^. The regulation of the *RNR4* gene may impact the production of malonyl-CoA through altering the availability of its precursors, but it could also be due to an improvement in growth by correctly balancing nucleotides. As with *BAP3*, a high-affinity broad-specificity amino acid permease^36^, where its null effect displays increased competitive fitness in minimal media^37^.

While in this study we used a gRNA library to increase fluxes towards malonyl-CoA, it should be noted that this is just one example of how this platform can be utilized, and other forms of selection such as the expression of gene libraries could also be incorporated^38–40^. Furthermore, this study utilized a limited gRNA library of only 168 genes, and in order to achieve higher accuracy and representation, future studies should consider utilizing a more comprehensive library^41,42^. Multiplexing strategies targeting a larger number of genes per cell could also be implemented^43^. Furthermore, other biosensors could be targeted for continuous enrichment of genotypes favoring the production of other targeted products, and could potentially be used in combination^44,45^. Finally, our approach for promoter selection relied on the strength of the promoter and its upstream position relative to malonyl-CoA in the metabolic pathway. However, it is possible that other metabolic genes might have been more optimal for this setup. To increase the sensitivity of the strain and further enhance the flux towards the targeted precursor, a multi-gene targeting strategy involving more than two promoters could be implemented.

In conclusion, our research sheds light on the innovative potential of using a growth-sensitive platform strain for the continuous evolution towards the production of valuable metabolites. Our study effectively demonstrates the viability of this approach and lays a foundation for future research and advancements in the field of microbial engineering.

## Authors’ contributions

RF and FD conceived the study and designed the experiments. KAH carried out the experiments. RF and KAH analyzed the data. RF and KAH wrote the manuscript. JN and FD supervised the study. All authors contributed to the analysis and the discussion of the results. All authors read and approved the final manuscript.

## Acknowledgments

The authors would like to thank Xiaowei Li for providing the mcr expression system, as well as Gustav Edman for initial discussions.

