## Supplementary Materials for "Biosensor-Assisted Laboratory Evolution of Malonyl-CoA production in *Saccharomyces cerevisiae*"

^5^ Current address: Department of Genetics, Harvard Medical School, Boston, Massachusetts, USA.

^6^ Current address: Novo Nordisk Foundation Center for Biosustainability, Technical University of Denmark, DK2800 Kgs. Lyngby, Denmark

^†^ Contributed equally

**^*^ Corresponding author**

**Supplementary figures**

- Figure S1. Growth performance of AHP strains.
- Figure S2. Growth performance of recovery from canavanine treated cells and high concentrated treated cells.
- Figure S3. Gradual gRNA enrichment.
- Figure S4. Gradual gRNA distribution enrichment for the ALE.
- Figure S5. Gradual gRNA distribution enrichment for the ALE without counter- selection.
- Figure S6. Mutation profiles of MPC01 with and without canavanine treatment.
- Figure S7. Characterization of the enriched gRNAs on growth fitness in MCP01C.

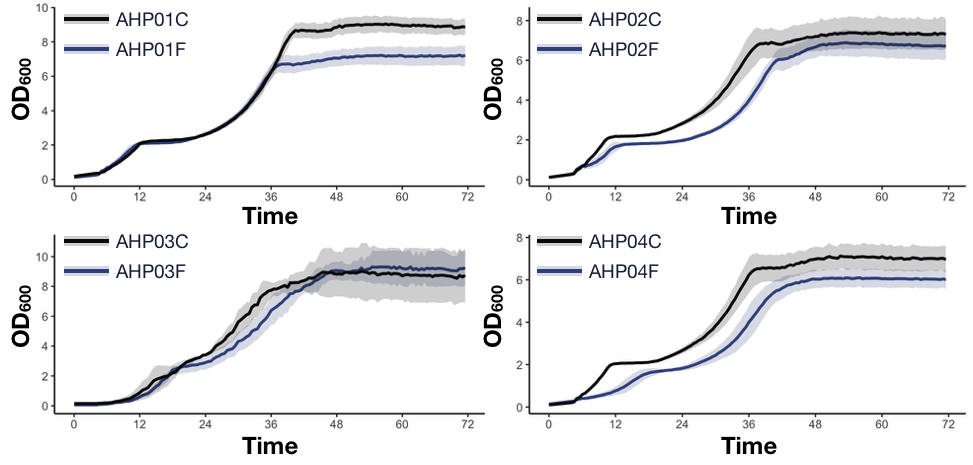

**Figure S1. Growth performance of AHP strains.** Growth curves of AHP strains with and without FapR (AHP01 = WT, AHP02 = P*_ACS2_*Δ::P*_TEF1_*BS123, AHP03 = P*_PGK1_*Δ::P*_TEF1_*BS123, AHP04 = P*_ACS2_*Δ::P*_TEF1_*BS123-P*_PGK1_*Δ::P*_TEF1_*BS123). Content shown as mean ± SD of three biological replicates grown in defined minimal medium with 2% glucose.

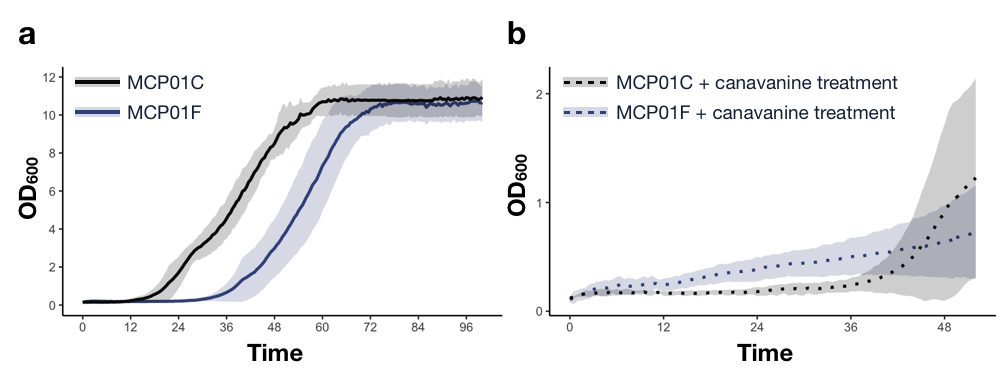

**Figure S2. Growth on canavanine treated cells. a.** Growth profile of the engineered platform strain MCP01 after treated with 3 ng·μL^-1^ canavanine. **b.** Growth curves of MCP01 in 10 ng·μL^-1^ canavanine medium. Content shown as mean ± SD of three biological replicates, grown in defined minimal medium with 2% glucose.

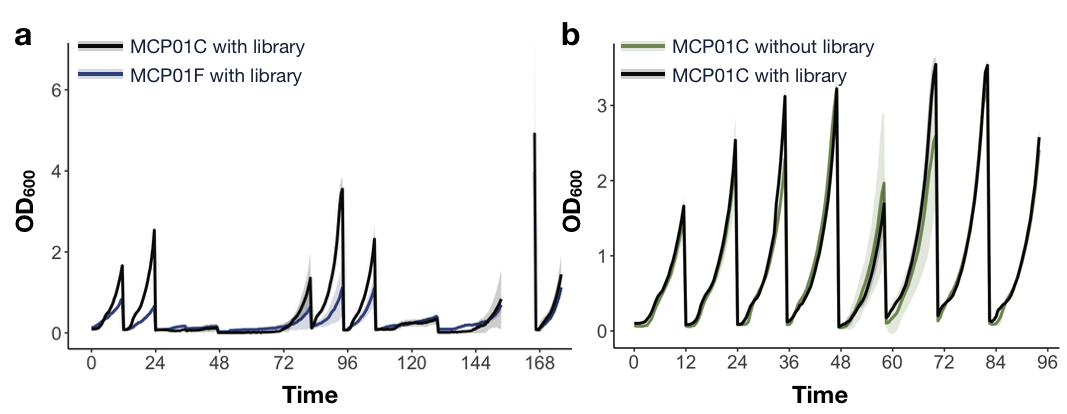

**Figure S3. ALE growth profiles in MPC01 strains. a.** Complete growth profile of the ALE. Diluted every 12 hours, after 24 hours and 108 hours, cells were exposed to canavanine treatment. No data were collected between 153 and 165 hours. **b.** Growth comparison of the engineered platform strain MCP01C with and without library. All cultivations were done in 2% minimal medium. Data shown as mean ± SD of three biological replicates.

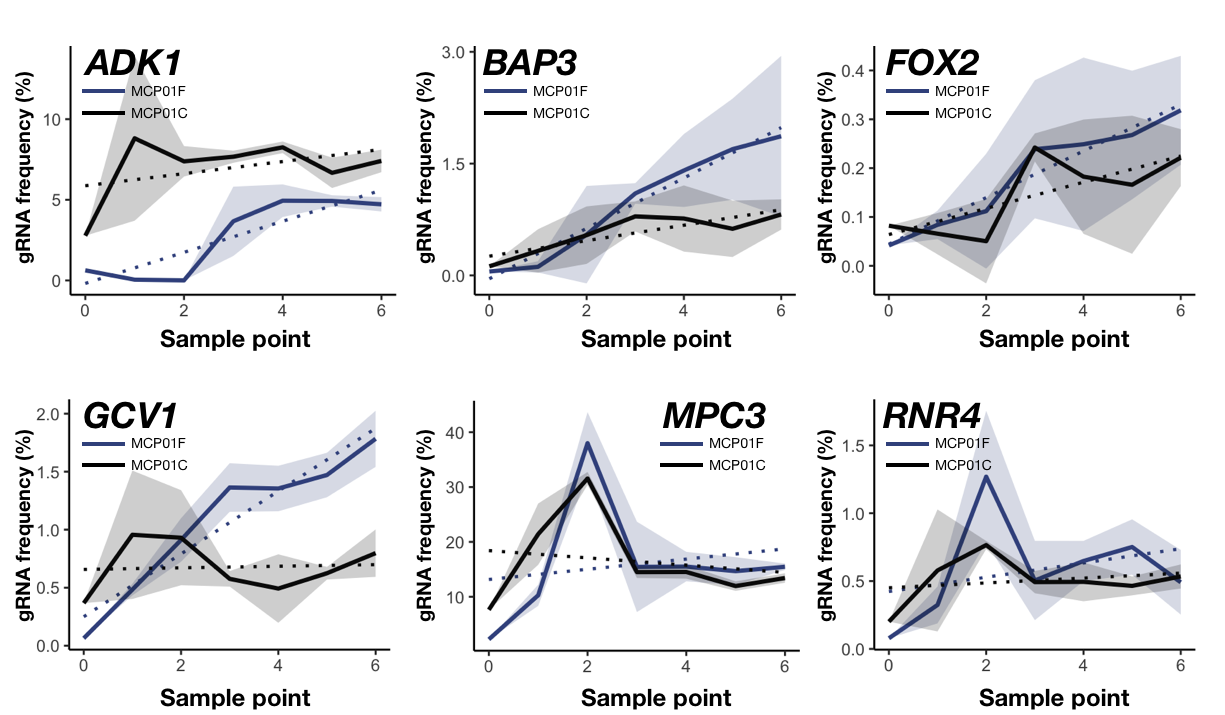

**Figure S4. Gradual gRNA distribution enrichment during the ALE.** The significant enriched gRNAs distribution under each sample point. Comparing gRNA expressed in MCP01F and MCP01C. Frequency shown as mean ± SD of three biological replicates, grown in defined minimal medium with 2% glucose. Dotted line represent a linear trendline.

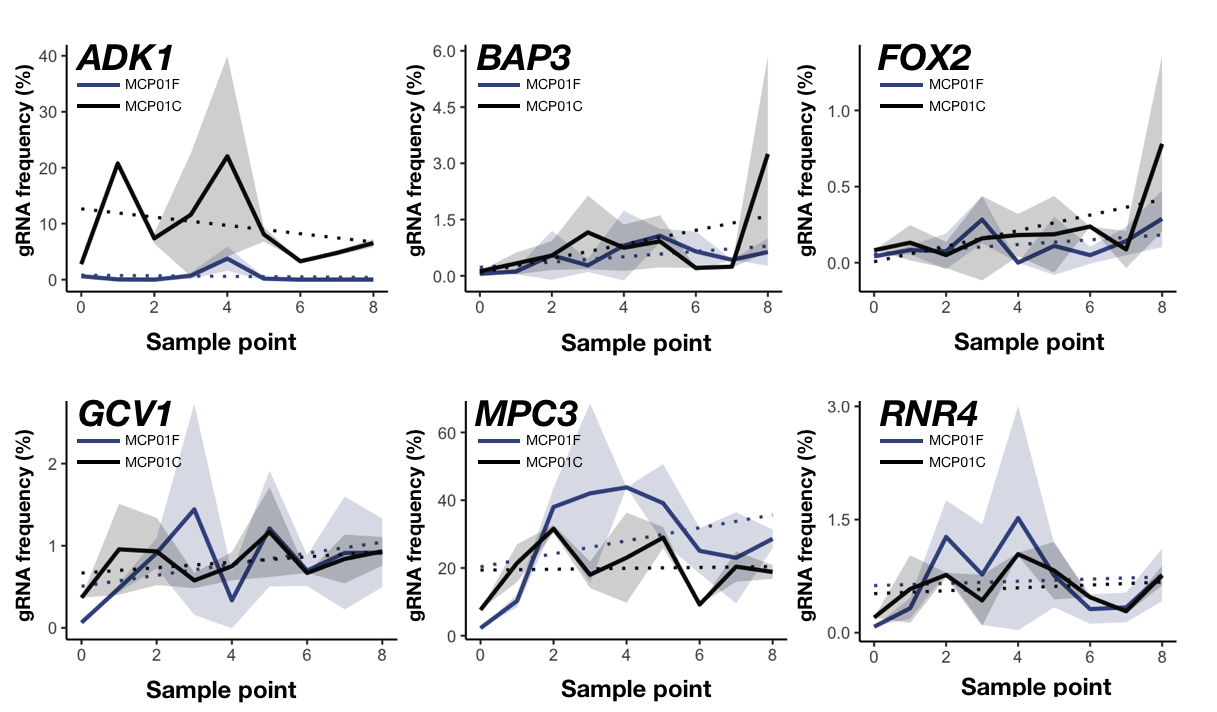

**Figure S5. Gradual gRNA distribution enrichment during the ALE without counter-selection treatment.** The significant enriched gRNAs distribution under each sample point. Comparing gRNA expressed in MCP01F and MCP01C. Frequency shown as mean ± SD of three biological replicates, grown in defined minimal medium with 2% glucose. Dotted lines represent a linear trendline.

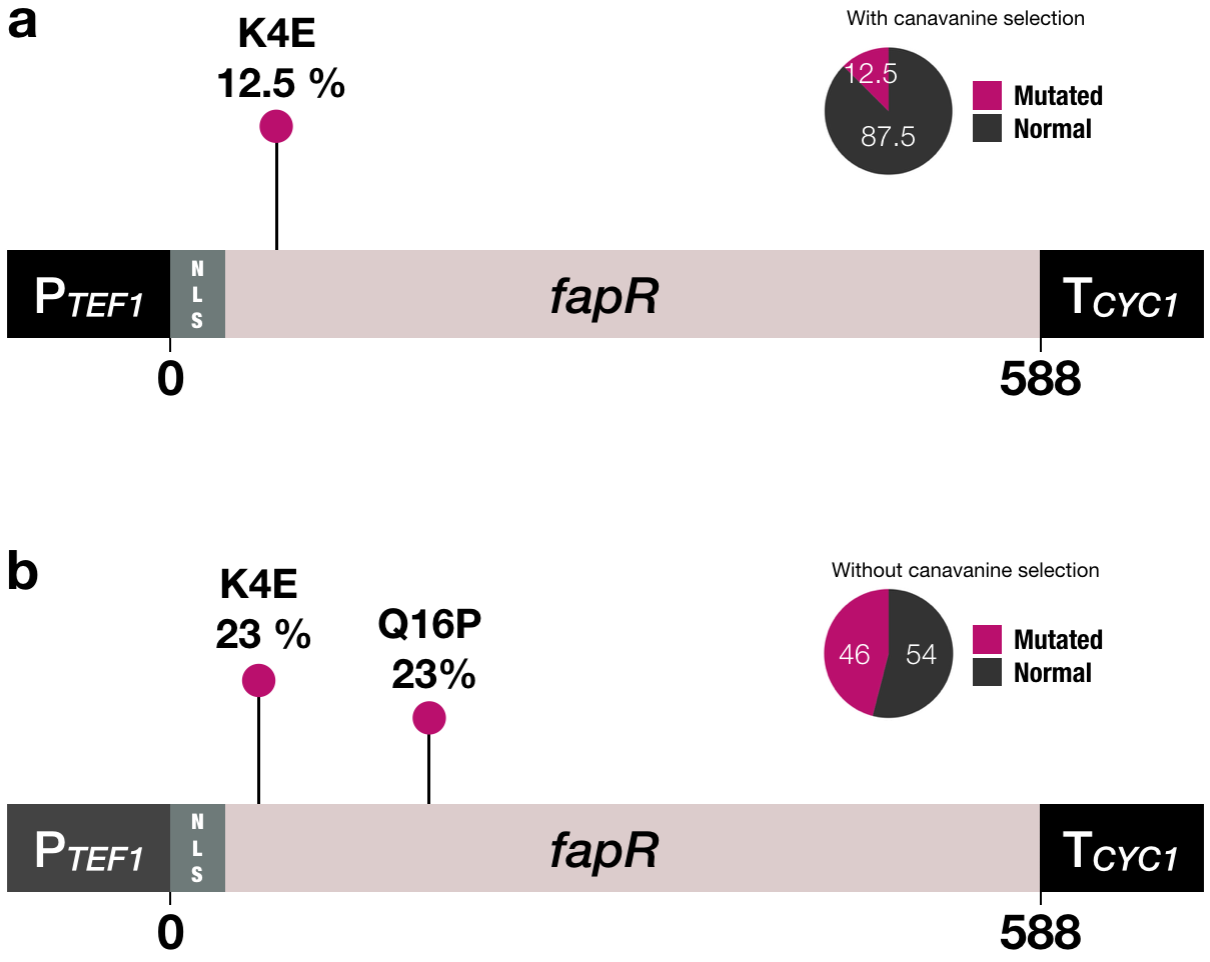

**Figure S6. Mutation profiles of MPC01 with and without canavanine treatment.** **a.** Mutation profile from 8 sequenced colonies after 176 hours (8 transferring) with canavanine treatment. One colony showed a mutation, K4E. **b.** Mutation profile from 13 sequenced colonies after 96 hours (8 transferring) without canavanine treatment. Six colonies showed a mutation, K4E or Q16P.

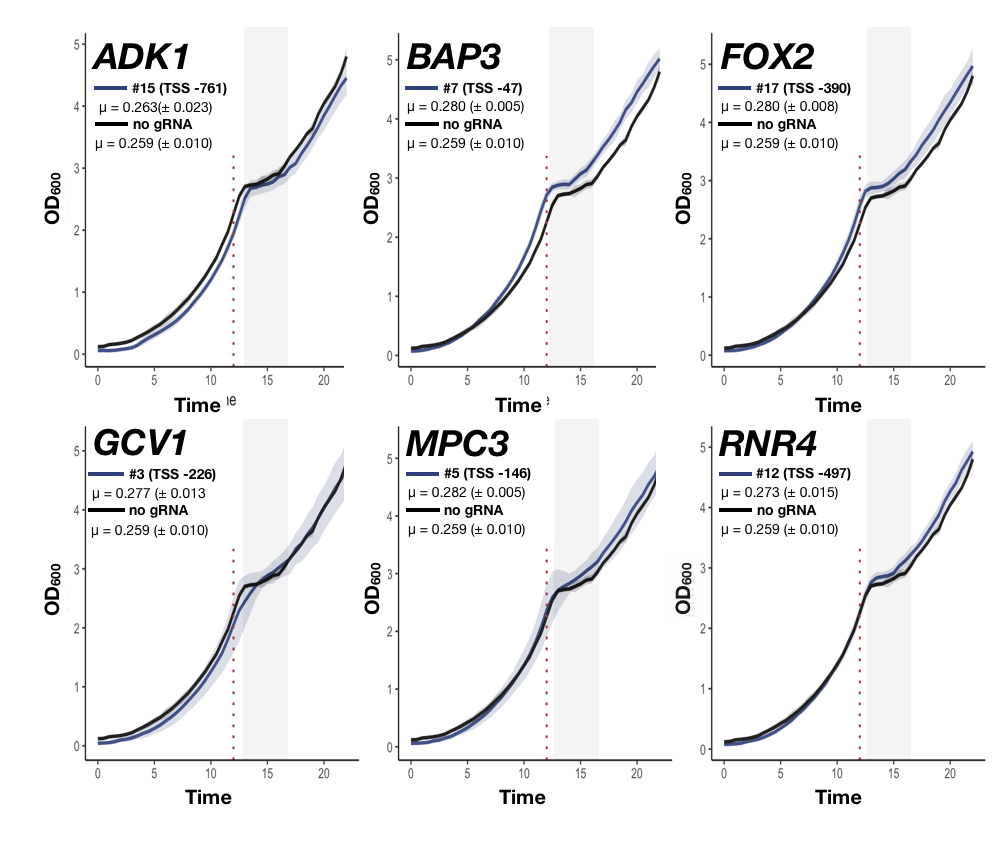
**Figure S7. Characterization of the enriched gRNAs on growth fitness in MCP01C.** Overall effect of expressing the enriched gRNAs in the control, MCP01C. The content shown as mean ± SD of three biological replicates and two technical replicates were monitored with a Growth Profiler 960. The growth rate was calculated between the start of the exponential phase to the 12-hour mark for all replicates ± SD. Outliers were removed (initial OD_600_ > 0.15).
